# Visualization of the Cdc48 AAA+ ATPase protein unfolding pathway

**DOI:** 10.1101/2023.05.13.540638

**Authors:** Ian Cooney, Heidi L. Schubert, Karina Cedeno, Hsien-Jung L. Lin, John C Price, Christopher P Hill, Peter S Shen

## Abstract

The Cdc48 AAA+ ATPase is an abundant and essential enzyme that unfolds substrates in multiple protein quality control pathways. The enzyme includes two conserved AAA+ ATPase cassettes, D1 and D2, that assemble as hexameric rings with D1 stacked above D2. Here, we report an ensemble of structures of Cdc48 affinity purified from lysate in complex with the adaptor Shp1 in the act of unfolding substrate. Our analysis reveals a continuum of structural snapshots that spans the entire translocation cycle. These data reveal new elements of Shp1-Cdc48 binding and support a “hand-over-hand” mechanism in which the sequential movement of individual subunits is closely coordinated. D1 hydrolyzes ATP and disengages from substrate prior to D2, while D2 rebinds ATP and re-engages with substrate prior to D1, thereby explaining the dominant role played by D2 in substrate translocation/unfolding.

## Introduction

The conserved AAA+ ATPase Cdc48 (aka. p97 or VCP) is an essential and abundant “segregase” that functions across many cellular pathways to separate proteins from various contexts, including organelle membranes, ribosomes, chromatin, and protein complexes. Interactions with substrate adaptors allows Cdc48 to function in multiple pathways, with involvement in specific pathways presumably underlying its importance in degenerative diseases and cancers^1,2^. Mutations in Cdc48 cause multisystem proteinopathy, a disease associated with the degeneration of the brain, bones, and muscles, although the underlying mechanism of disease progression is poorly understood. Additionally, Cdc48 is upregulated in various types of cancer, and specific inhibitors of Cdc48 are currently under evaluation in clinical trials.

Cdc48 subunits comprise an N-terminal (N) domain followed by tandem AAA+ ATPase cassettes, D1 and D2, which are the motors that bind and hydrolyze ATP to drive substrate translocation/unfolding. D1 and D2 each comprise small and large domains, and share sequence and structural similarity. Multiple structures show that Cdc48 assembles as a homo-hexamer of stacked D1 and D2 rings, with each of the 12 cassettes contributing conserved pore loop 1 (PL1) and pore loop 2 (PL2) residues that bind the unfolded substrate in a groove through the central hexameric pore^3–6^. The N-domain sits atop the complex and is the primary binding site of substrate-recruiting adaptors.

A wide variety of protein substrates are brought to Cdc48 by adaptors, which can also regulate subcellular localization and ATP hydrolysis rates. One of the most abundant Cdc48 adaptors in budding yeast is Shp1, which regulates protein phosphatase-1 (PP1) complex assembly and stability^7,8^. The human orthologs of Shp1, p47 and p37, function in post-mitotic Golgi reassembly and PP1 activation, respectively^9,10^. In human cells, the PP1 catalytic subunit is held in an inactive state through interactions with its binding partners SDS22 and Inhibitor-3 (I3), and is activated by the Cdc48-mediated unfolding of I3 in a p37-dependent manner. The unfolding of I3 frees PP1 to bind downstream activating cofactors and allows it to perform its critical role in transcriptional regulation as a serine/threonine phosphatase^7^. The relationship between Cdc48 and PP1 is presumably conserved from yeast to human because the yeast orthologs of the PP1 complex are enriched in native pulldowns of Shp1 (human, PP1-SDS22-I3; yeast, Glc7-Sds22-Ypi1)^3^.

Insights into how AAA+ ATPases drive substrate unfolding have been derived from substrate-bound structures of Cdc48 and related AAA+ unfoldases^3–6,11–26^. The leading model envisions unfolding driven by translocation of the substrate in an extended conformation through the hexamer pore by a “hand-over-hand” (aka. “conveyer belt”) mechanism^3,5,21,27^. In this model, D1 and D2 form an ATP-stabilized helical spiral that presents a peptide-binding groove with the optimal symmetry to bind substrate in an extended β-strand conformation, with an array of equivalent dipeptide binding sites formed at the interfaces of adjacent Cdc48 subunits. Typically, five hexamer subunits adopt the spiraling configuration and bind substrate, while the other subunit appears to be transitioning between the ends of the spiral. Progression along the unfolding/translocation cycle results from ATP hydrolysis-mediated dissociation of a Cdc48 subunit from the lagging end of the spiral while ATP binding-mediates association at the leading end of the spiral to bind the next dipeptide of the substrate. Although we strongly favor this hand-over-hand mechanism, we note that alternatives have been suggested^28–31^.

Here, we determined an ensemble of cryo-EM structures of the Cdc48-Shp1 complex in the act of processing substrate. The structures belong to a series of nine sequential snapshots that represent a continuum of subunit movements during substrate translocation. The motion of each Cdc48 subunit is directly linked to its neighboring subunits through inter-subunit interactions. These structures provide a complete view of the entire protein translocation cycle and rationalize a model in which the binding and hydrolysis of ATP and the engagement and disengagement of substrate are highly coordinated.

## Results and Discussion

### Overall structure

Native Cdc48-Shp1 complexes were purified by co-immunoprecipitation of FLAG-tagged Shp1 from budding yeast lysates, as described previously^3,32^. Active, substrate-bound complexes were enriched by stabilizing the particles with the ATP analog ADP•BeF_x_, which traps Cdc48 in the process of unfolding endogenous substrate. Cryo-EM images of the purified particles were recorded, and subsequent data processing produced an overall consensus charge density map at 2.9 Å resolution (Supplemental Fig. 1). The reconstruction closely resembled our previously reported 3.7 Å and 3.6 Å resolution structures of yeast^3^ and human^5^ Cdc48, respectively, in which the D1 hexamer is stacked on top of D2 in a right-handed helical arrangement for subunits A-E, with A being the topmost subunit that is engaged with substrate in both D1 and D2, while subunit F is disengaged from the helical array and the substrate.

### 3D variability analysis reveals the Cdc48 translocation pathway continuum

In order to visualize conformational heterogeneity in Cdc48, we applied 3D variability analysis (3DVA)^33^ to the 2.9 Å resolution dataset. This revealed a component that tracked the continuous motion of subunit F between the bottom and the top of the helical stack (Movies 1, 2). Particles were split into ten classes along this component, nine of which were resolved to global resolutions between 3.2 Å to 4.3 Å (Supplemental Figs. 2, 3). The classes were assigned based on the relative position of subunit F along the translocation pathway: Class 1 displays subunit F in its lowest position relative to the helical array, and class 9 displays subunit F at the top of the helical array, re-engaged with substrate and subunit A in D2 but still disengaged in D1 (Movie 3). The intermediate classes 2-8 display subunit F along a sequential trajectory between classes 1 and 9.

Models for subunits A, B, and C were closely superimposable across all classes. Subunit D displayed some relative motion within the small ATPase domain, but was always part of the AAA+ helix and engaged with substrate. We define the D1 and D2 domains as being engaged if the alpha carbon atom (Cα) of the PL1 residue M288 (D1) or W561 (D2) is within 6 Å of a substrate Cα; in contrast, all disengaged subunits had distances of at least 9 Å. Subunit E exhibited positional variability and disengagement from the AAA+ helix and substrate across some, but not all, of the classes while, as expected, subunit F is always disengaged in D1 and is in a different position in all classes. This indicates that more than one subunit becomes disengaged from substrate at distinct stages along the translocation trajectory. The consensus class overlays best with classes 1-3, which have the largest number of contributing particles and show subunit E engaged with substrate and subunit F in an intermediate position along the translocation trajectory^3^. In order to map the complete path taken by one subunit as it transitions from the bottom to the top of the AAA+ helix, the states of subunit E were mapped onto the path taken by subunit F by overlap of subunits A and B onto the positions of subunits B and C (Fig. 1).

**Figure 1.**
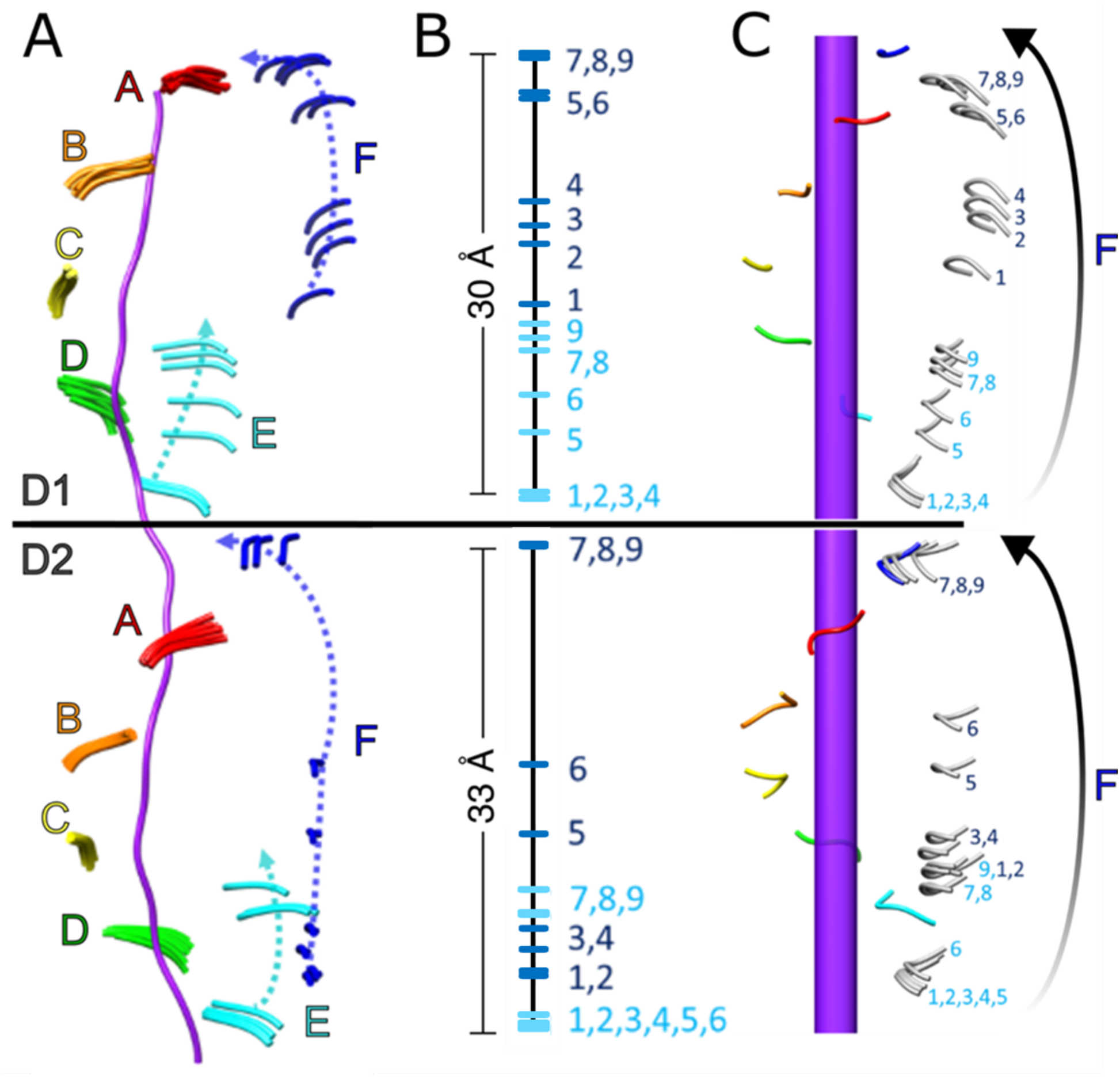
Trajectory of pore loop-1 (PL1) motion during substrate translocation. (A) Overlay of all nine models of PL1 residues (ribbons) wrapped around substrate (purple). (B) Position of PL1 models from subunits E (cyan) and F (blue) relative to substrate. (C) Positioning of subunit E PL1 models along the path of subunit F reveals the continuous trajectory of the substrate-disengaged subunit from the bottom to the top of each helical stack. Substrate axis represented as purple rod. The start and end positions of the transitioning subunit inferred from extrapolation of the helical assembly are colored blue, with intermediate states inferred from class 1-9 conformations of subunits E and F shown in grey.

### D2 engages with substrate over a larger fraction of the translocation cycle than D1

Our analysis revealed that the trajectories for D1 and D2 are slightly out of phase at the start and end of the translation pathway (Fig. 1, Movie 3). As subunit F progresses toward the top of the helical stack, subunit E disengages from the bottom and follows the translocation path of subunit F. At the start of the translocation trajectory, D1 of subunit E is engaged with the substrate in classes 1-4 and becomes disengaged in class 5, while D2 of subunit E does not disengage from substrate until class 7. This indicates that D1 disengages from the substrate before D2. In contrast, at the end of the trajectory, D1 of subunit F is still disengaged from substrate in class 9, while D2 has re-engaged to form a canonical substrate binding pocket with the corresponding pore loop residues from subunit A. In essence, D2 of subunit F has already assumed the position of subunit A in this class while D1 is still moving toward that position.

Thus, D2 engages the substrate before D1. Our finding that D2 spends more time than D1 engaged with the substrate is consistent with better resolved substrate density in D2 than in D1 (Supplemental Fig. 4). Presumably, D2 binds substrate more tightly because it contains conserved large hydrophobic residues at critical positions in PL1 (W561 and Y562), while D1 displays more flexible and smaller residues at the equivalent positions (M288 and A289). These observations rationalize how D2 plays the dominant role in imparting the translocation force that drives unfolding by gripping onto substrate more tightly^34,35^. This is also consistent with the finding that catalytically inactivating mutants in D1 retain some substrate processing activity while the corresponding mutations in D2 completely abrogate translocation^34^.

### Small and large ATPase domain motions are asynchronous and linked to movement of adjacent subunits

The large and small ATPase domains of subunit D exhibit variable local resolutions and were modeled as separate rigid bodies as they begin to move behind subunit E. The large domains of subunit D remain essentially stationary as subunit E disengages from the substrate and moves between classes 5-9. In contrast, the small domain of subunit D undergoes an ∼8° rotation in both D1 and D2 (Fig. 2A). The large and small domains of subunit E were also fitted as separate rigid bodies as they trail behind subunit F. Although the D1 and D2 large domains and the D2 small domain remain stationary, the D1 small domain undergoes a ∼5° rotation between classes 1-4 before the large domain of subunit E has disengaged from substrate (Fig. 2B).

**Figure 2.**
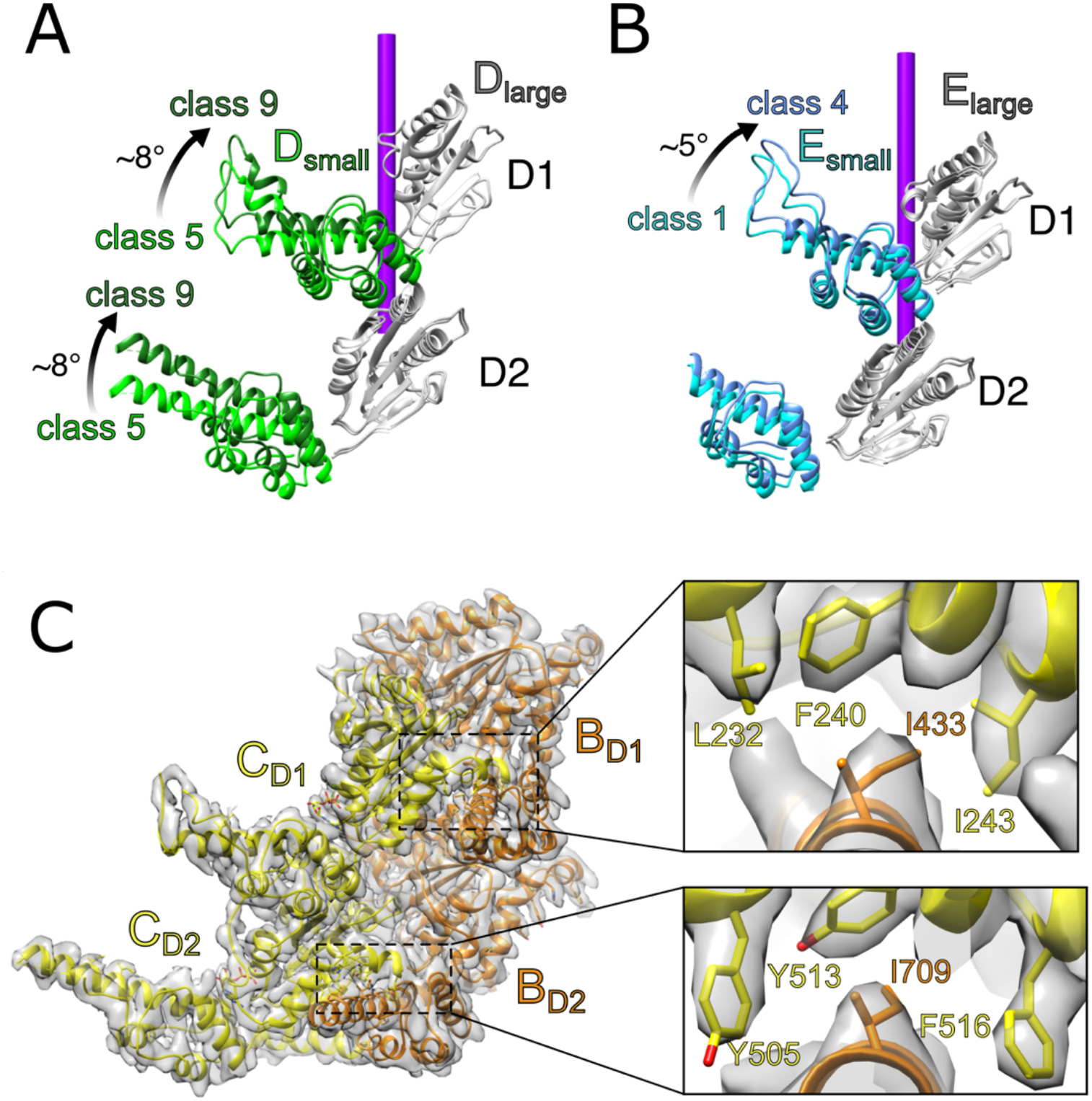
Flexible motions within individual subunits during substrate translocation. (A) The small domains of subunit D in both D1 and D2 move independently from their large domains between classes 5-9. (B) The small domain of subunit E from D1 is relatively mobile between classes 1-4 while its large domain and D2 domains remain stationary. (C) Inter-subunit contacts between neighboring large and small ATPase domains (BC interface shown). The large domains form hydrophobic binding pockets comprising F240, I243, and L232 of D1 and Y505, Y513, and F516 of D2. A conserved isoleucine (D1, I433; D2, I709) is inserted into each binding pocket from the neighboring subunit. This inter-subunit interaction is likely preserved during subunit movement.

The relative motions of small and large domains within a subunit appear to be driven by inter-subunit contacts between the small domain of one subunit and the large domain of the following subunit, which are largely preserved as subunits move along the translocation pathway (Fig. 2C). As clearly defined for subunits B and C in the center of the helical stack, the primary interaction for D1 involves I422 of the B subunit small domain inserting into a hydrophobic pocket formed by L232, F240, and I243 of the C subunit large domain. For D2, contacts include an analogous interaction occurs between subunit B small domain I709 and subunit C large domain Y505, Y513, and F516. These interactions appear to be maintained throughout the entire translocation pathway, although the poor local resolution in mobile subunits does not allow definitive visualization in all cases^12,15^. Thus, the leading subunit’s large domain appears to drag the small domain of the following subunit, linking the movement of adjacent subunits. A similar interaction was described in maintaining the interface of neighboring large and small domains of Vps4, suggesting that these inter-subunit contacts may be a conserved feature of AAA+ enzymes^11^.

The 9 conformations span the entire substrate translocation cycle and support the hand-over-hand mechanism in which Cdc48 “walks” along the substrate to pull unfolded substrate through the central pore. Previous structures of substrate-bound Cdc48 report that a single Cdc48 subunit of the homohexamer disengages from substrate during the “hand-over-hand” cycle^3,4^. Here, we find that at some stages of the reaction cycle, two subunits are fully disengaged (Fig. 1, Movie 3), and three subunits (D, E, and F) display intermediate conformations that are linked though maintenance of the small domain-large domain interactions of adjacent subunits.

### Nucleotide states are coupled to subunit movement

Our reconstructions allow assignment of nucleotide density as corresponding to ATP, ADP, or apo (Supplemental Table 1). In all classes, the D1 and D2 active sites of subunits A, B, and C display nucleotide density corresponding to ATP (ADP•BeF_x_) coordinated with Mg^2+^ (Supplemental Fig. 5). ATP density is also present in early classes of subunit D (classes 1-5 in D1 and classes 1-6 in D2) as well as in later classes of subunit F (classes 8-9 in D2) (Supplemental Figs. 6, 7). Walker B glutamates activate a water molecule for nucleophilic attack on the gamma phosphate of ATP (D1, E315; D2, E588). The sensitivity of negatively charged residues to electron irradiation leads to degradation of density surrounding the carboxylate side chains^36^, thereby limiting our ability to model the precise positions of the catalytic glutamate side chain atoms directly, although in all active sites displaying ADP•BeF_x_ density, the Walker B glutamate main chain and nucleotide superimposed closely with other well-defined structures of AAA+ ATPase-ATP complexes^3,5^. This is consistent with the possibility that all of the ADP•BeF_x_ (ATP)-bound states in classes 1-9 are competent to catalyze ATP hydrolysis.

Arginine finger residues in D1 (R369, R372) and D2 (R645, R648) are provided by the following subunit to coordinate nucleotide phosphates and promote ATP hydrolysis by stabilizing the pentavalent double negatively charged transition state^37^. Their side chains are well defined for active sites that bind ADP•BeF_x_ in the consensus structure, with the guanidium group within hydrogen-bonding distance to a gamma phosphate oxygen atom (i.e., an F in BeF_x_) (Supplementary Fig. 5). For active sites that bind ADP•BeF_x_ in lower resolution classes but lack resolution to visualize the arginine side chain, the arginine main chain and bound nucleotide are superimposable with the high-resolution structures, consistent with guanidinium-phosphate coordination. As with the Walker B geometry, this is consistent with all of the ATP (ADP•BeF_x_)-bound active sites adopting a catalytically competent conformation.

The transition in density from ATP to ADP in the later classes of subunit D (class 6 in D1 and class 7 in D2) indicates that ATP hydrolysis occurs in this subunit at the D-E interface (Supplementary Fig. 6). Given the superimposable, apparently active conformation at all of the ATP-binding active sites, it is not clear why hydrolysis should be favored at this specific location. One possibility is suggested by the motion of the subunit E large domain along the translocation pathway, which may result in a structurally modest but catalytically critical repositioning of the finger arginine to allow catalysis, as has been suggested for Vps4^15^ (Supplementary Fig. 8). Another possibility for preferential ATP hydrolysis at subunit D is that the highly constrained environment at the subunit A, B, and C active sites creates an unfavorable environment for phosphate release after ATP hydrolysis. In contrast to the AB, BC and CD interfaces, the nucleotide-binding pocket at the subunit DE interface active site becomes widened as subunit E progresses along the translocation path (from class 6 in D1 and from class 7 in D2), which may provide a favorable environment for phosphate release. This is reflected in the increased distance as the arginine fingers from subunit E move away from the nucleotide at the subunit D active site in class 6 for D1 and in class 7 for D2 (Supplemental Fig. 8).

Nucleotide density at the subunit E active site appears as ADP in early classes and is no longer observed (i.e., becomes empty/apo, or too low resolution to discern) in classes 5-9 in D1 and classes 7-9 in D2 (Table 1, Supplementary Fig. 7). Likewise, subunit F nucleotide density is variable, being absent in all classes for D1 and in classes 1-7 for D2, but corresponding to ADP•BeF_x_•Mg^2+^ in D2 of classes 8-9 as it approaches subunit A and begins to reconnect to the Cdc48 helical array.

The ensemble of classes visualized establishes the relationship between the nucleotide hydrolysis cycle and substrate translocation (Fig. 3A). Subunits A-D are tightly engaged with substrate and their inter-subunit interfaces (AB, BC, and CD) are also compact in ATP-bound states (Fig. 3, Table 1). A progression from substrate-engaged to disengaged state is observed for subunit E, which also correlates with a transition of nucleotide state from ATP to ADP at the DE interface (subunit D active site). This transition may be a result of a widening of the DE interface as subunit E and its nucleotide-coordinating arginine fingers move along the translocation trajectory away from subunit D, thereby promoting ATP hydrolysis and phosphate release at the DE interface. Opening of the active site as subunit E moves along the translocation pathway opens the nucleotide binding site to allow exchange of ADP to apo to ATP. At the end of the translocation cycle, subunit F re-engages substrate and binds ATP and subunit A at the top of the helical stack. This subunit therefore becomes the “new” subunit A and the translocation cycle advances by one subunit.

**Figure 3.**
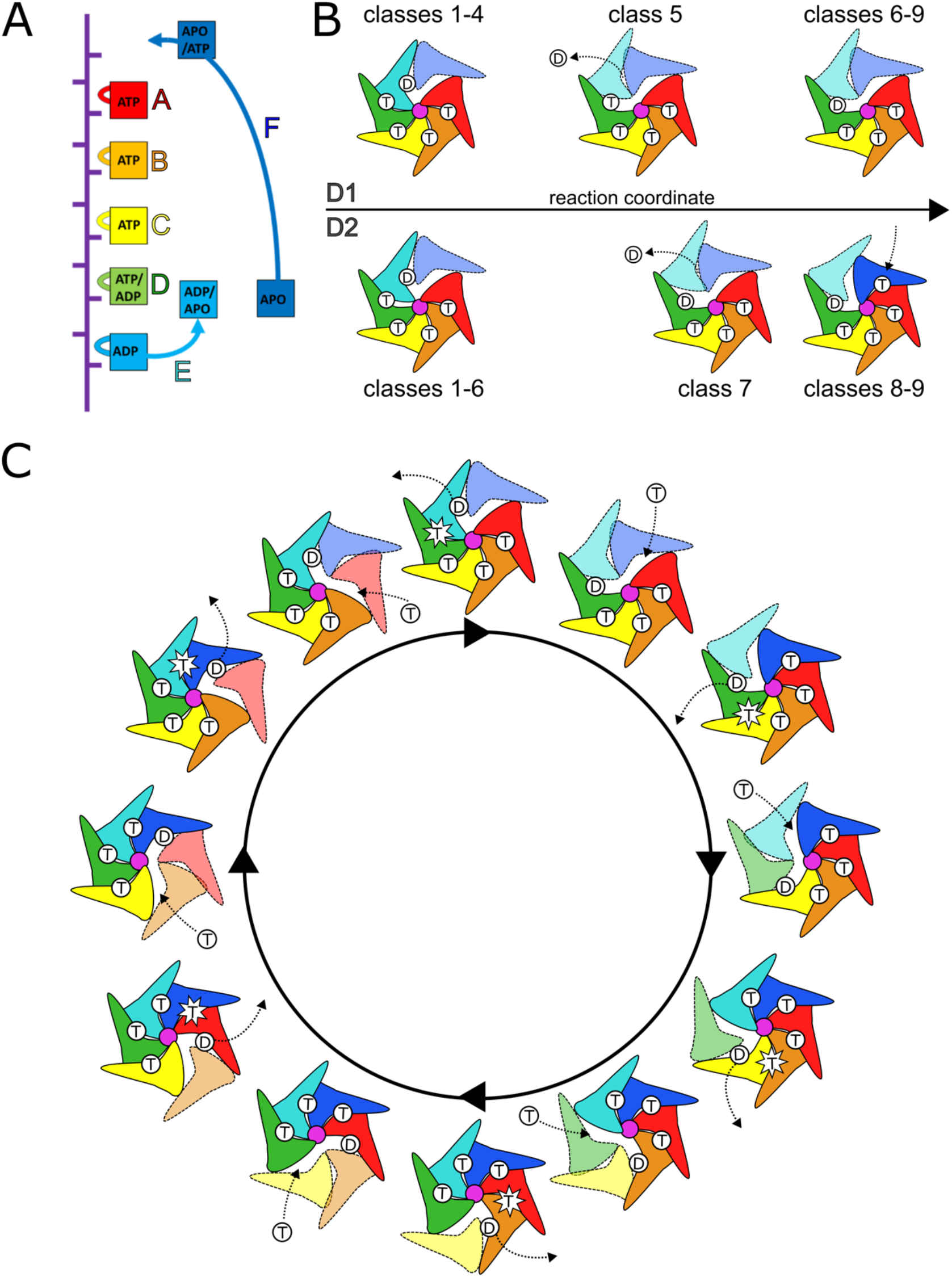
Cdc48 translocation model. (A) Nucleotide state assignments of individual Cdc48 subunits. (B) D1 and D2 do not move in lockstep with each other. D1 hydrolyzes ATP before D2. D2 binds ATP before D1. Substrate-disengaged subunits shown in lighter shades and separated from substrate (purple). (C) General translocation cycle of the Cdc48 hexamer. Each subunit undergoes sequential steps of ATP hydrolysis, disengagement from substrate, ATP binding, and re-engagement to substrate. Note that this is applicable to both D1 and D2, though the rings are not always in phase with each other. T, ATP; D, ADP.

### Native Cdc48-Shp1 complexes are enriched with PP1 and ubiquitylated substrate

Our previous work^3^ indicated that native yeast Cdc48-Shp1 purifications were enriched with components of the protein phosphatase-1 (PP1) complex, which includes Glc7, Sds22, and Ypi1. These results were consistent with studies of human proteins indicating that the orthologous p97-p37 complex unfolds human Ypi1 (Inhibitor-3, I3) to allow the formation of PP1 holoenzymes^7,10^. This prompted us to investigate if PP1 components are a substrate of the Cdc48-Shp1 complex in yeast. To test this, we immobilized native Shp1-associated complexes from budding yeast lysates using ADP•BeF_x_ to trap bound substrates. Samples were then eluted with excess ATP to outcompete ADP•BeF_x_ and release substrates from the immobilized resin (Supplementary Fig. 9). Mass spectrometry proteomics of the ATP eluate revealed an enrichment of PP1 components compared to controls. Diglycine ubiquitin signatures were not detected on PP1 peptides, suggesting that ubiquitin-independent unfolding of Ypi1/I3 is conserved from yeast to human. However, ubiquitin (Ubi4) was enriched in the ATP elution sample, indicating that the Cdc48-Shp1 complex also processes ubiquitylated substrates that are distinct from PP1. An enrichment of ubiquitin was also observed in our recent characterization of the homologous human p97-p47 complex^5^. Interestingly, p47 does not appear to be associated with PP1^38^, which suggests that yeast Shp1 acts on both ubiquitin-dependent and independent substrates while human p37 and p47 have specialized roles that have diverged from the ancestral Shp1.

### Substrate and Shp1 contacts with the Cdc48 N domains

The Cdc48 N-terminal domains are associated with low-resolution density in the consensus reconstruction, consistent with their known conformational variability ^39^. We therefore performed focused classification to probe for additional features associated with these domains. This revealed a sub-class (class P) that contained a large density attached to the N-domain of subunit A (Fig. 4A, Supplemental Fig. 10) into which the crystal structure of the human PP1-SDS22 (Glc7-Sds22) subcomplex fit as a rigid body^40^. The subunit B N-domain was associated with density corresponding to the Shp1 UBX domain, as previously reported ^3,5,41^. This structure is consistent with a recent report of reconstituted human PP1 complex bound to p97 and p37 in which density for SDS22 and PP1 were observed on an N-domain of p97 of one subunit and p37 UBX density on an N-domain of a following subunit^42^. Focused classification over other subunits (B-F) did not resolve PP1 density, probably because these classes contained too few particles. Ypi1 is not apparent within the modeled density, though it may be part of unassignable low-resolution density observed above the central pore that connects to the substrate modeled in the translocation pore (Supplemental Fig. 10).

**Figure 4.**
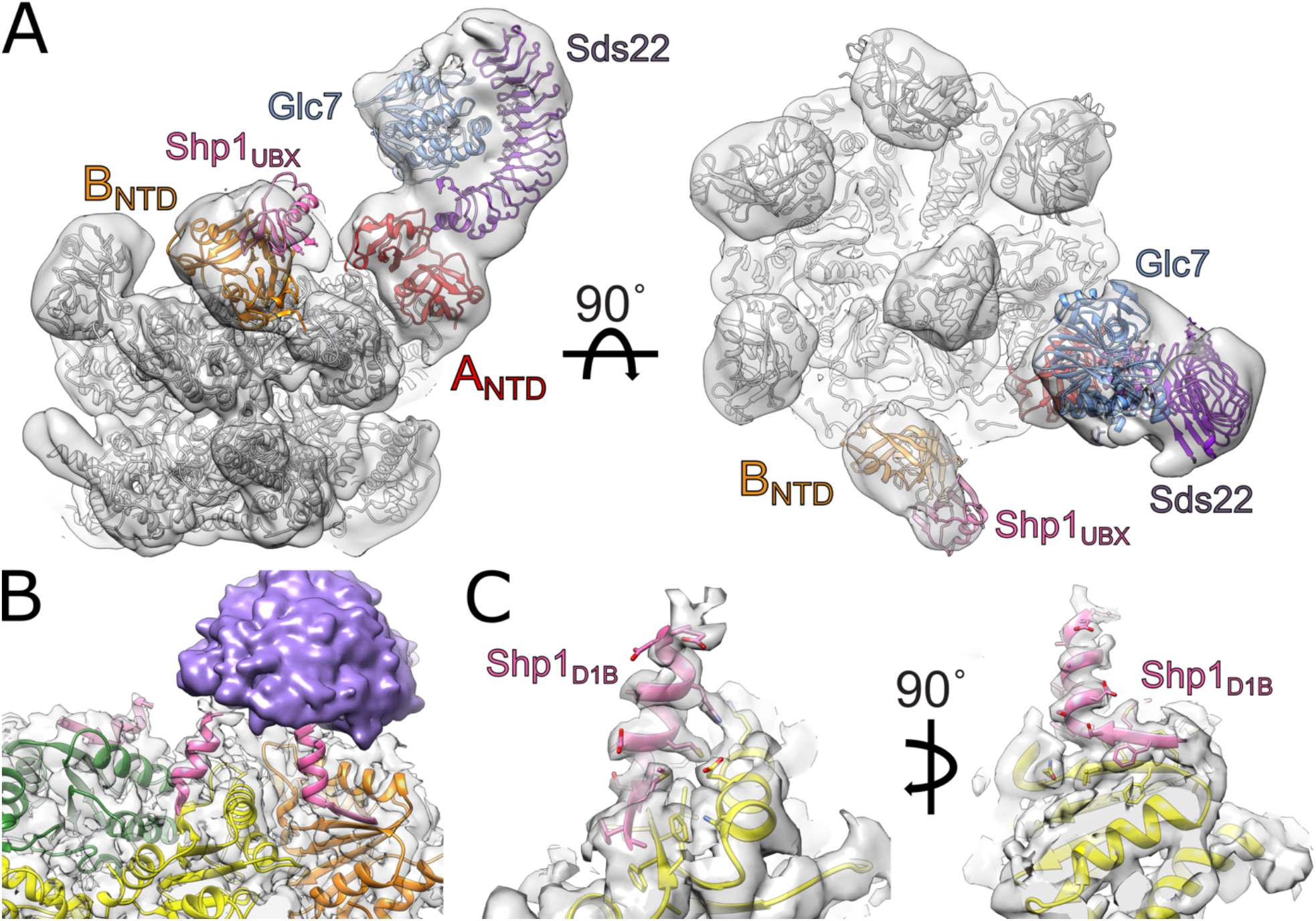
Substrate and adaptor densities associated with Cdc48. (A) Rigid-body fitting of PP1 (Glc7-Sds22, PDB 6OBN) and UBX (PDB 1S3S) crystal structures into Cdc48 reconstruction. (B) Model and density of Shp1 D1-binding (D1B) motif bound to D1 large AAA+ ATPase domains of subunits B, C, and D (pink). The D1B density is connected to unstructured density above the central pore (purple). (C) Close-up model and density of Shp1 D1B motif (pink) bound to D1 of subunit C (yellow).

### Shp1 contacts the D1 large ATPase domain

The D1 large ATPase domain includes a solvent-exposed β-sheet that is well resolved in subunits B, C, and D (Fig. 4B). In the consensus structure, these subunits display additional density that is associated with the outermost β-strand (residues 276-279). The density over subunit C was of sufficient quality to build 14 additional residues that include a four-residue β-strand that extends the β-sheet, followed by ten additional α-helical residues directed toward the extra density above the central pore of the hexamer (Fig. 4C).

To determine the source of this density, we employed ColabFold^43^ to predict potential interactions between the D1 large domain and Shp1 as well as other possible adaptors and substrates^5^. The best scoring model showed a specific region of Shp1 (residues 125-138) that fit precisely as a rigid-body within the extra density (Supplemental Figure 11). Densities at subunits B and D are consistent with the model built at subunit C, and it seems probable that equivalent structures are present at subunits A, E, and F but are not apparent due to the poorer local resolution for those subunits. This additional interaction between this D1-binding motif (D1B) of Shp1 and D1 likely strengthens adaptor binding by complementing binding of the Shp1 UBX domain with the Cdc48 NTD. It may also promote optimal positioning of the Shp1-bound Ypi1 substrate towards the central pore of the hexamer.

## Conclusion

Our structural analysis revealed how the Cdc48 translocation pathway is driven by ATP hydrolysis, phosphate release, ADP dissociation, and ATP binding, and refines earlier models by showing that D1 and D2 do not move in lockstep (Fig. 3B). Specifically, D1 hydrolyzes ATP, releases phosphate and dissociates from substrate before D2, while D2 rebinds ATP and re-engages with substrate before D1. This occurs sequentially with adjacent subunit nucleotide states and positions being linked by inter-subunit contacts and up to three subunits observed moving along the translocation pathway simultaneously. The PP1 complex was seen to contact the Cdc48 N-domain and a new motif of Shp1 was observed bound to D1 in interactions that were only observed in a subset of the least mobile subunits but are likely distributed with partial occupancy over all subunits. Our findings provide a foundation to investigate conservation of the translocation mechanism in other AAA+ ATPases and further investigate function and interactions of Shp1, and can inform efforts to develop novel therapeutics such as inhibitors of Cdc48/p97.

## Acknowledgements

This work was supported by grants to PSS (NIH R35 GM133772), CPH (NIH U54 AI170856 and R01 GM112080), JCP (NIH R01 AG066874), and IC (NIH F31 CA254427). We thank the University of Utah Arnold and Mabel Beckman Center for Cryo-EM, Center for High Performance Computing, and the Fritz B. Burns Biological Mass Spectrometry Facility at Brigham Young University for support in cryo-EM, computation, and mass spectrometry, respectively.

## Methods

### Protein purification

Cdc48-Shp1 complexes were purified as described previously^3,32^. In brief, a C-terminal 3xFLAG tag was introduced to the endogenous locus of the Shp1 gene in budding yeast (strain BY4741). Yeast cells were grown to mid-log phase, harvested, lysed, and clarified supernatant was used as the basis of “lysate-to-grid” purification and cryo-EM specimen preparation.

### Cryo-EM specimen preparation

Crosslinking was performed with 0.1% glutaraldehyde for 10 minutes and quenched with excess Tris to a final concentration of ∼30 mM. Specimens where prepared on UltraAuFoil R1.2/1.3 Au300 mesh grids and were glow discharged for 60 seconds at 25 mA with a Pelco easiGlow unit. 3.5 μl of sample were applied to the grid, followed by blotting and vitrification using a Mk. II Vitrobot with a blot time of 5 seconds into liquid ethane.

### Data collection and image reconstruction

Cryo-EM movies were recorded on a 300 kV Titan Krios G3 with a K3 direct detector using SerialEM^44^ with a dose rate of 19.3 e/pix/sec at a nominal magnification of 81,000x (corresponding to a super-resolution pixel size of 0.54 Å/pixel). Each movie contained 40 frames recorded over a span of 2.0 seconds each, corresponding to a total electron exposure of ∼45 e/Å^2^. A total of 5,913 movie were recorded. Patch motion correction and CTF estimation were performed in cryoSPARC and micrographs were 2x binned to their physical pixel size of 1.08 Å/pixel ^45,46^.

### Cryo-EM data processing

All data processing was performed in CryoSPARC ^45^. Templates were generated using multiple rounds of 2D classification for particles identified by blob picking from a small subset of the movies collected (249 movies). These templates were used in template picking of the entire data set. This led to selection of 4,690,491 particles, which were reduced to 1,456,150 particles after multiple rounds of 2D classification. Particles were subsequently re-extracted with a box size of 288 pixels from 5,847 movies, and further rounds of 2D classification resulted in 539,957 particles. Ab initio classification followed by heterogeneous refinement in CryoSPARC led to the sorting of 325,377 particles into a well-resolved class. Non-uniform refinement of these particles resulted in a 2.8 Å reconstruction of the Cdc48 complex (Supplemental Fig. 1).

### 3DVA analysis

3DVA analysis^33^ of the particles contributing to the 2.8 Å consensus structure revealed a component that displayed variability of mobile subunits, and was used in a 3DVA Display job to cluster the particles into 10 classes, of which 9 could be reconstructed using non-uniform refinement, with the 10^th^ class failing to reconstruct due to limited particle number. Inspection of the 9 reconstructed classes revealed asymmetric substrate-bound structures with variable positioning of mobile subunits. The mobile subunits had limited local resolution, so another round of 3DVA and 3DVA display was performed on all classes. After non-uniform refinement of the resulting clusters, resolution improvements were observed in 6 classes by excluding additional particles (Supplemental Fig. 2). This refinement led to final resolutions ranging between 3.2-4.3 Å for all classes (Supplemental Figs 2, 3, 12).

Particles were re-extracted with a larger box size of 360 pixels to include additional volume for substrate and adaptor interactions with the N-domain. 3DVA analysis of particles was then performed using a large mask over the N-domain and part of D1 (Supplemental Fig. 20), and particles containing density on the N-termini of subunit A were selected from clustering in 3DVA display. This was followed by multiple rounds of 2D classification, ab initio refinement, heterogeneous refinement, and further 3DVA analysis to isolate particles that contained this density. This led to 7,727 particles containing extra density, with non-uniform refinement yielding a reconstruction at 4.2 Å resolution.

### Model building, refinement, and validation

Model building was performed for each of the class reconstructions. Models of the D1 and D2 ATPase cassettes were built and subject to atomic refinement using Phenix 1.2.1^47^ for all A, B, C & D subunits, while the large and small domains of subunits E in classes 2-9 and F in classes 1-8 were fit as rigid bodies (from PDB 6OPC^3^) using Chimera^48^ and Coot v0.9.8.1 ^49^.

Substrate was built as described^3^. For subunits with ADP•BeF_x_ modeled, magnesium was restrained to the OG1 of the final Thr in the Walker A domain at a distance of 2.038 Å. To model pore loops for subunit F structures that lacked density, we used the well-defined structures of subunit A. The same approach was taken for subunit E pore loops using PDB 6OPC^3^. Rigid body fitting of Sds22/PP1 used PDB 60BN^50^ in UCSF Chimera for Sds22/PP1 density. All figures and movies where prepared in UCSF Chimera^48^. A summary of model refinement statistics is provided in Supplemental Table 2.

### ColabFold Structure Prediction

An initial search for interactors that would bind the beta strand of the D1 large domain was performed by performing predictions over ∼100 residue sections of Glc7, Sds22, Ypi1, Doa1, and Shp1 and investigating the interactor prediction for these residues against the D1 large domain of Cdc48. After finding an interaction between Shp1 and Cdc48, final predictions were made in ColabFold using sequences for residues 125-139 of Shp1 and the Cdc48 the D1 large domain (residues 219-379)^43^. Template mode was selected as pdb70 and 24 recycles were performed. This prediction fit well as a rigid body into the density for subunit C (Supplemental Fig. 11) of the consensus structure, and was used as a starting point for further refinement in Phenix.

### Mass spectrometry

Mass spectrometry methods were as described previously with minor adaptations^3^. Co-IP eluates from Shp1-FLAG in the presence of 1 mM ADP•BeF_x_ or 5 mM ATP, to elute putative substrate, were processed separately for proteomics analysis (n=3 for each sample). For each sample, snap-frozen eluates were thawed with the addition of 100 μL of 6 M guanidinium/HCl in 100 mM Tris/HCl pH 8.5 containing protease inhibitor cocktail (Sigma). Samples were washed with 6 M guanidine/HCl 100 mM Tris/HCl, pH 8.5 using centrifugal filters (30 kD MWCO). Cysteines were reduced using 10 mM TCEP and alkylated using 30 mM chloroacetamide. The guanidine solution was removed using two 25 mM ammonium bicarbonate (pH 8) washes. Proteins were resuspended in 25 mM ammonium bicarbonate (pH 8) and digested overnight at 37°C with 0.2 μg of MS-Grade trypsin (Pierce). Trypsin reactions were quenched using 1 mM PMSF, and trypsin was removed by centrifugal filtration. Samples were placed in mass spec vials, dried using a vacuum concentrator, and then resuspended at 1 μg/μL in 3% acetonitrile, 0.1% formic acid. Mass spectrometry data were collected using an Orbitrap Fusion Lumos mass spectrometer (Thermo Fisher Scientific) coupled to Ultimate 3000 liquid chromatography (LC) pump (Thermo Fisher Scientific). Peptides were separated using an EASY-Spray PepMap RSLC C18 column (Thermo Fisher Scientific). The mobile phase consisted of buffer A (0.1% formic acid in optima water) and buffer B (optima water and 0.1% formic acid in 80% acetonitrile). The peptides were eluted at 300 nL/min with the following gradients over 2 h: 3– 25% B for 80 min; 25–35% B for 20 min; 35–45% B for 8 min; 45–85% B for 2 min and 85% for 8 min. Data were acquired using the top speed method (3 s cycle). A full scan mass spectrum at resolution of 120,000 at 200 m/z mass was acquired in the Orbitrap with a target value of 4e5 and a maximum injection time of 50 ms. Peptides with charge states of 2-6 were selected from the top abundant peaks by the quadrupole for collisional dissociation (CID with normalized energy 30) MS/MS, and the fragment ions were detected in the linear ion trap with target AGC value of 1e4, a maximum injection time of 35 ms, and a dynamic exclusion time was set at 60 s. Precursor ions with ambiguous charge states were not fragmented. PEAKS Studio software (version 10) was used for de novo sequencing and database searching to identify proteins in the raw MS data, and to quantify, filter (quality-control), and normalize the quantitation data for each protein^51^. Peptides were identified from MS/MS spectra by searching against the Swiss-Prot Saccharomyces cerevisiae (strain ATCC 204508 / S288c) database (downloaded August 2020) with a reverse sequence decoy concatenated database^52^. Variables for the search were as follows: enzyme was set as trypsin with one missed cleavage site. Carbamidomethylation of cysteine was set as a fixed modification while N-terminal acetylation and methionine oxidation were set as variable modifications. A false positive rate of 0.01 was set as the maximum for peptides and proteins. The minimum length of peptide was set to 7 amino acids. At least 2 peptides were required for protein identification. The precursor mass error of 20 ppm was set for the precursor mass, and the mass error was set as 0.3 Da for the MSMS. Label-free quantitation was enabled with MS1 tolerance ±20 ppm and a MS2 tolerance ±50 ppm. The probability of each protein being a Shp1 interactor was calculated by comparing 3 replicates of each co-IP using the SAINT software package^53^.

## Notes

### Competing Interest Statement

The authors have declared no competing interest.

## References

1. Johnson M.A. et al. The Cure VCP Scientific Conference 2021: Molecular and clinical insights into neurodegeneration and myopathy linked to multisystem proteinopathy-1 (MSP-1). Neurobiol Dis. 169, (2022).

2. Anderson, D. J. et al. Targeting the AAA ATPase p97 as an Approach to Treat Cancer through Disruption of Protein Homeostasis. Cancer Cell 28, 653–665 (2015).

3. Cooney, I. et al. Structure of the Cdc48 segregase in the act of unfolding an authentic substrate. Science (1979) 365, 502–505 (2019).

4. Twomey, E. C. et al. Substrate processing by the Cdc48 ATPase complex is initiated by ubiquitin unfolding. Science (1979) 365, (2019).

5. Xu, Y. et al. Active conformation of the p97-p47 unfoldase complex. Nat Commun 13, 2640 (2022).

6. Pan, M. et al. Mechanistic insight into substrate processing and allosteric inhibition of human p97. Nat Struct Mol Biol 28, 614–625 (2021).

7. Weith, M. et al. Ubiquitin-Independent Disassembly by a p97 AAA-ATPase Complex Drives PP1 Holoenzyme Formation. Mol Cell 72, 766–777.e6 (2018).

8. Cheng, Y.-L. & Chen, R.-H. Assembly and quality control of the protein phosphatase 1 holoenzyme involves the Cdc48–Shp1 chaperone. J Cell Sci 128, 1180–1192 (2015).

9. Uchiyama, K. p97/p47-Mediated Biogenesis of Golgi and ER. J Biochem 137, 115–119 (2005).

10. van den Boom, J. et al. Targeted substrate loop insertion by VCP/p97 during PP1 complex disassembly. Nat Struct Mol Biol 28, 964–971 (2021).

11. Han, H., Monroe, N., Sundquist, W. I., Shen, P. S. & Hill, C. P. The AAA ATPase Vps4 binds ESCRT-III substrates through a repeating array of dipeptide-binding pockets. Elife 6, (2017).

12. Su, M. et al. Mechanism of Vps4 hexamer function revealed by cryo-EM. Sci Adv 3, (2017).

13. Wang, L., Myasnikov, A., Pan, X. & Walter, P. Structure of the AAA protein Msp1 reveals mechanism of mislocalized membrane protein extraction. Elife 9, (2020).

14. Zehr, E. A., Szyk, A., Szczesna, E. & Roll-Mecak, A. Katanin Grips the β-Tubulin Tail through an Electropositive Double Spiral to Sever Microtubules. Dev Cell 52, 118–131.e6 (2020).

15. Monroe, N., Han, H., Shen, P. S., Sundquist, W. I. & Hill, C. P. Structural basis of protein translocation by the Vps4-Vta1 AAA ATPase. Elife 6, (2017).

16. Han, H., Monroe, N., Sundquist, W. I., Shen, P. S. & Hill, C. P. The AAA ATPase Vps4 binds ESCRT-III substrates through a repeating array of dipeptide-binding pockets. Elife 6, (2017).

17. Han, H. et al. Structure of Vps4 with circular peptides and implications for translocation of two polypeptide chains by AAA+ ATPases. Elife 8, (2019).

18. Puchades, C. et al. Structure of the mitochondrial inner membrane AAA+ protease YME1 gives insight into substrate processing. Science (1979) 358, eaao0464 (2017).

19. de la Peña, A. H., Goodall, E. A., Gates, S. N., Lander, G. C. & Martin, A. Substrate-engaged 26 S proteasome structures reveal mechanisms for ATP-hydrolysis–driven translocation. Science (1979) 362, eaav0725 (2018).

20. Dong, Y. et al. ryo-EM structures and dynamics of substrate-engaged human 26S proteasome. Nature 565, 49–55 (2019).

21. Ripstein, Z. A., Huang, R., Augustyniak, R., Kay, L. E. & Rubinstein, J. L. Structure of a AAA+ unfoldase in the process of unfolding substrate. Elife 6, (2017).

22. Gates, S. N. et al. Ratchet-like polypeptide translocation mechanism of the AAA+ disaggregase Hsp104. Science (1979) 357, 273–279 (2017).

23. Deville, C. et al. Structural pathway of regulated substrate transfer and threading through an Hsp100 disaggregase. Sci Adv 3, (2017).

24. Yu, H. et al. ATP hydrolysis-coupled peptide translocation mechanism of Mycobacterium tuberculosis ClpB. Proceedings of the National Academy of Sciences 115, (2018).

25. White, K. I., Zhao, M., Choi, U. B., Pfuetzner, R. A. & Brunger, A. T. Structural principles of SNARE complex recognition by the AAA+ protein NSF. Elife 7, (2018).

26. Cho, C. et al. Structural basis of nucleosome assembly by the Abo1 AAA+ ATPase histone chaperone. Nat Commun 10, 5764 (2019).

27. Han, H. et al. Structure of spastin bound to a glutamate-rich peptide implies a hand-over-hand mechanism of substrate translocation. Journal of Biological Chemistry 295, 435–443 (2020).

28. Fei, X. et al. Structures of the ATP-fueled ClpXP proteolytic machine bound to protein substrate. Elife 9, (2020).

29. Sauer, R. T., Fei, X., Bell, T. A. & Baker, T. A. Structure and function of ClpXP, a AAA+ proteolytic machine powered by probabilistic ATP hydrolysis. Crit Rev Biochem Mol Biol 57, 188–204 (2022).

30. Cordova, J. C. et al. Stochastic but Highly Coordinated Protein Unfolding and Translocation by the ClpXP Proteolytic Machine. Cell 158, 647–658 (2014).

31. Fei, X. et al. Structures of the ATP-fueled ClpXP proteolytic machine bound to protein substrate. Elife 9, (2020).

32. Cooney, I. et al. Lysate-to-grid: Rapid Isolation of Native Complexes from Budding Yeast for Cryo-EM Imaging. Bio Protoc 13, (2023).

33. Punjani, A. & Fleet, D. J. 3D variability analysis: Resolving continuous flexibility and discrete heterogeneity from single particle cryo-EM. J Struct Biol 213, 107702 (2021).

34. Bodnar, N. O. & Rapoport, T. A. Molecular Mechanism of Substrate Processing by the Cdc48 ATPase Complex. Cell 169, 722–735.e9 (2017).

35. Esaki, M., Islam, Md. T., Tani, N. & Ogura, T. Deviation of the typical AAA substrate-threading pore prevents fatal protein degradation in yeast Cdc48. Sci Rep 7, 5475 (2017).

36. Bartesaghi, A., Matthies, D., Banerjee, S., Merk, A. & Subramaniam, S. Structure of β-galactosidase at 3.2-Å resolution obtained by cryo-electron microscopy. Proceedings of the National Academy of Sciences 111, 11709–11714 (2014).

37. Nadanaciva, S., Weber, J., Wilke-Mounts, S. & Senior, A. E. Importance of F 1 -ATPase Residue α-Arg-376 for Catalytic Transition State Stabilization. Biochemistry 38, 15493–15499 (1999).

38. Kracht, M. et al. Protein Phosphatase-1 Complex Disassembly by p97 is Initiated through Multivalent Recognition of Catalytic and Regulatory Subunits by the p97 SEP-domain Adapters. J Mol Biol 432, 6061–6074 (2020).

39. Banerjee, S. et al. 2.3 Å resolution cryo-EM structure of human p97 and mechanism of allosteric inhibition. Science (1979) 351, 871–875 (2016).

40. Choy, M. S. et al. SDS22 selectively recognizes and traps metal-deficient inactive PP1. Proceedings of the National Academy of Sciences 116, 20472–20481 (2019).

41. Dreveny, I. et al. Structural basis of the interaction between the AAA ATPase p97/VCP and its adaptor protein p47. EMBO J 23, 1030–1039 (2004).

42. van den Boom, J., Meyer, H. & Saibil, H. Structural basis of ubiquitin-independent PP1 complex disassembly by p97. bioRxiv 2022.06.24.497491 (2022) doi:10.1101/2022.06.24.497491.

43. Mirdita, M. et al. ColabFold: making protein folding accessible to all. Nat Methods 19, 679–682 (2022).

44. Mastronarde, D. N. Automated electron microscope tomography using robust prediction of specimen movements. J Struct Biol 152, 36–51 (2005).

45. Punjani, A., Rubinstein, J. L., Fleet, D. J. & Brubaker, M. A. cryoSPARC: algorithms for rapid unsupervised cryo-EM structure determination. Nat Methods 14, 290–296 (2017).

46. Rohou, A. & Grigorieff, N. CTFFIND4: Fast and accurate defocus estimation from electron micrographs. J Struct Biol 192, 216–221 (2015).

47. Afonine, P. V. et al. Real-space refinement in PHENIX for cryo-EM and crystallography. Acta Crystallogr D Struct Biol 74, 531–544 (2018).

48. Pettersen, E. F. et al. UCSF Chimera: A visualization system for exploratory research and analysis. J Comput Chem 25, 1605–1612 (2004).

49. Emsley, P., Lohkamp, B., Scott, W. G. & Cowtan, K. Features and development of Coot. Acta Crystallogr D Biol Crystallogr 66, 486–501 (2010).

50. Choy, M. S. et al. SDS22 selectively recognizes and traps metal-deficient inactive PP1. Proceedings of the National Academy of Sciences 116, 20472–20481 (2019).

51. Zhang, J. et al. PEAKS DB: De Novo Sequencing Assisted Database Search for Sensitive and Accurate Peptide Identification. Molecular & Cellular Proteomics 11, M111.010587 (2012).

52. Bairoch, A. The SWISS-PROT protein sequence database and its supplement TrEMBL in 2000. Nucleic Acids Res 28, 45–48 (2000).

53. Teo, G. et al. SAINTexpress: Improvements and additional features in Significance Analysis of INTeractome software. J Proteomics 100, 37–43 (2014).

