## Supplemental Figures for "Visualization of the Cdc48 AAA+ ATPase protein unfolding pathway"

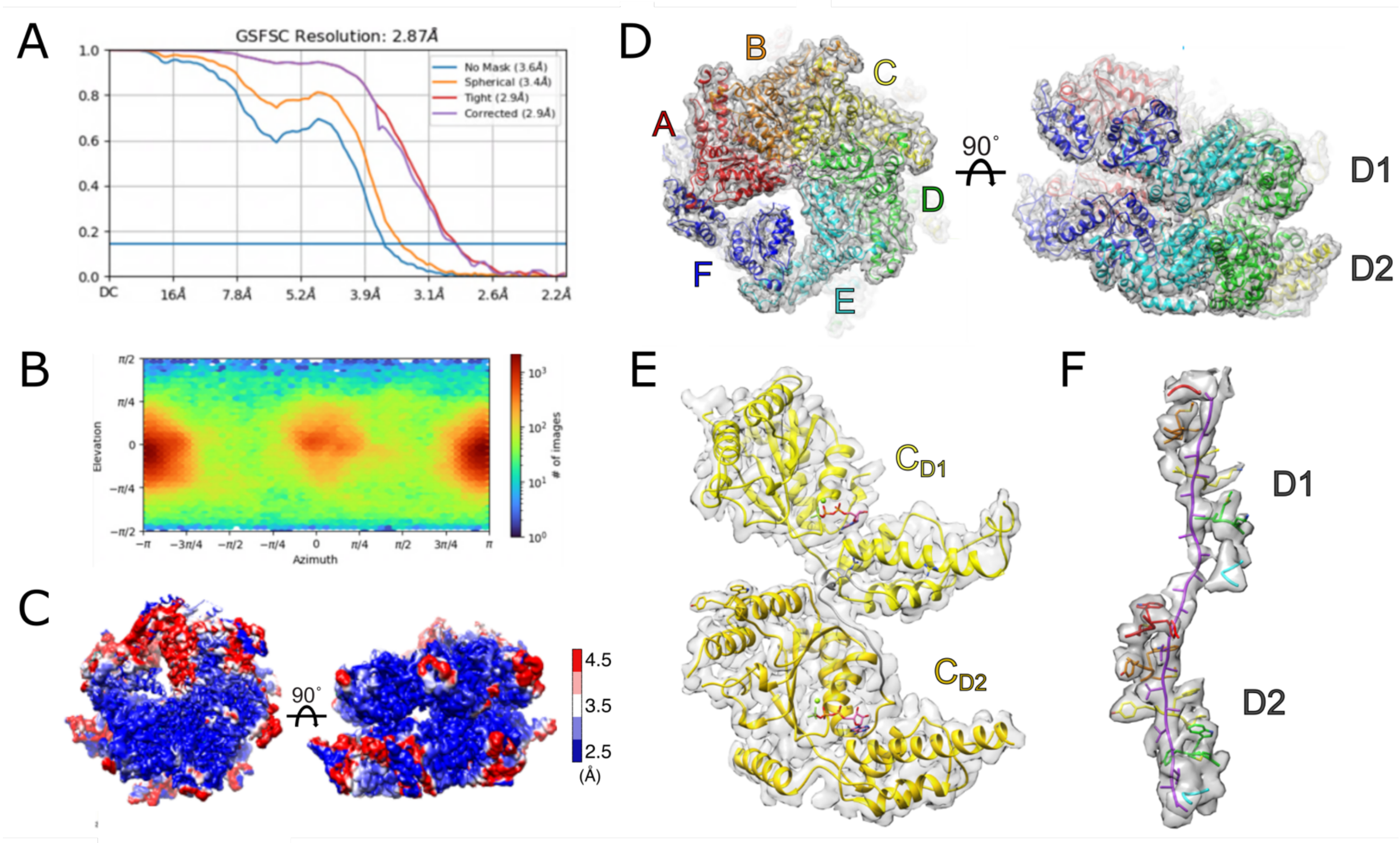


**Supplemental Figure 1. Consensus structure of Cdc48 bound to substrate.** (A) Gold standard FSC plot of 2.9 Å consensus reconstruction. (B) Orientation distribution plot. (C) Local resolution heat map. (D) Consensus map density is zoned around displayed residues to a distance of 2.5 Å at 3.5σ. (E) Model and zoned density at 6.5σ of subunit C. (F) Density of pore loop interactions with the substrate displayed at 4σ.

A

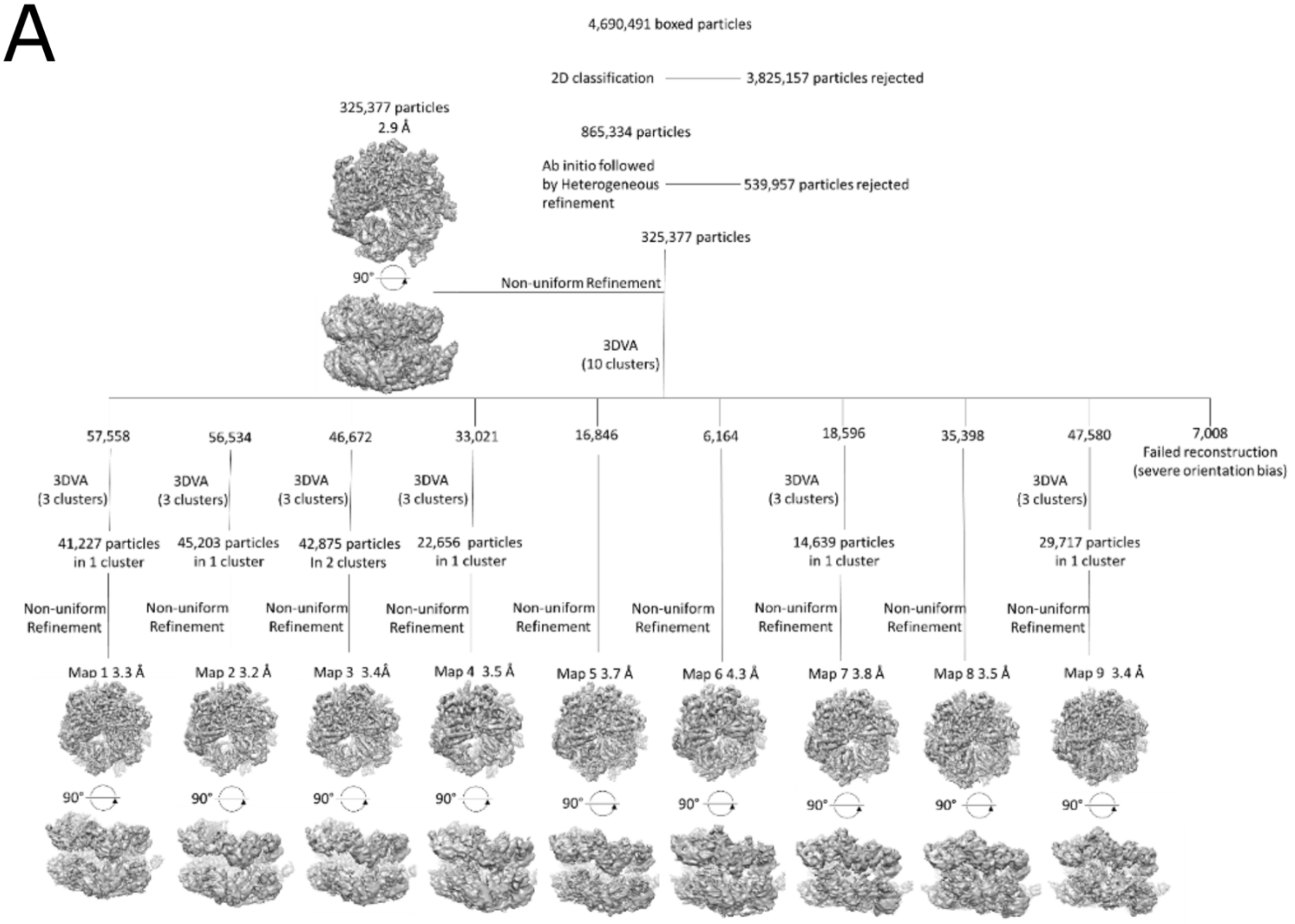


B

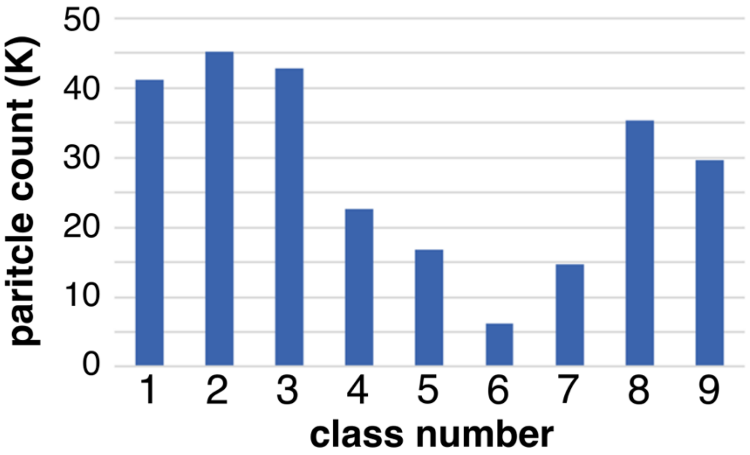


**Supplementary Figure 2. Cryo-EM image processing**. (A) Image processing workflow. (B) Particle number distribution.

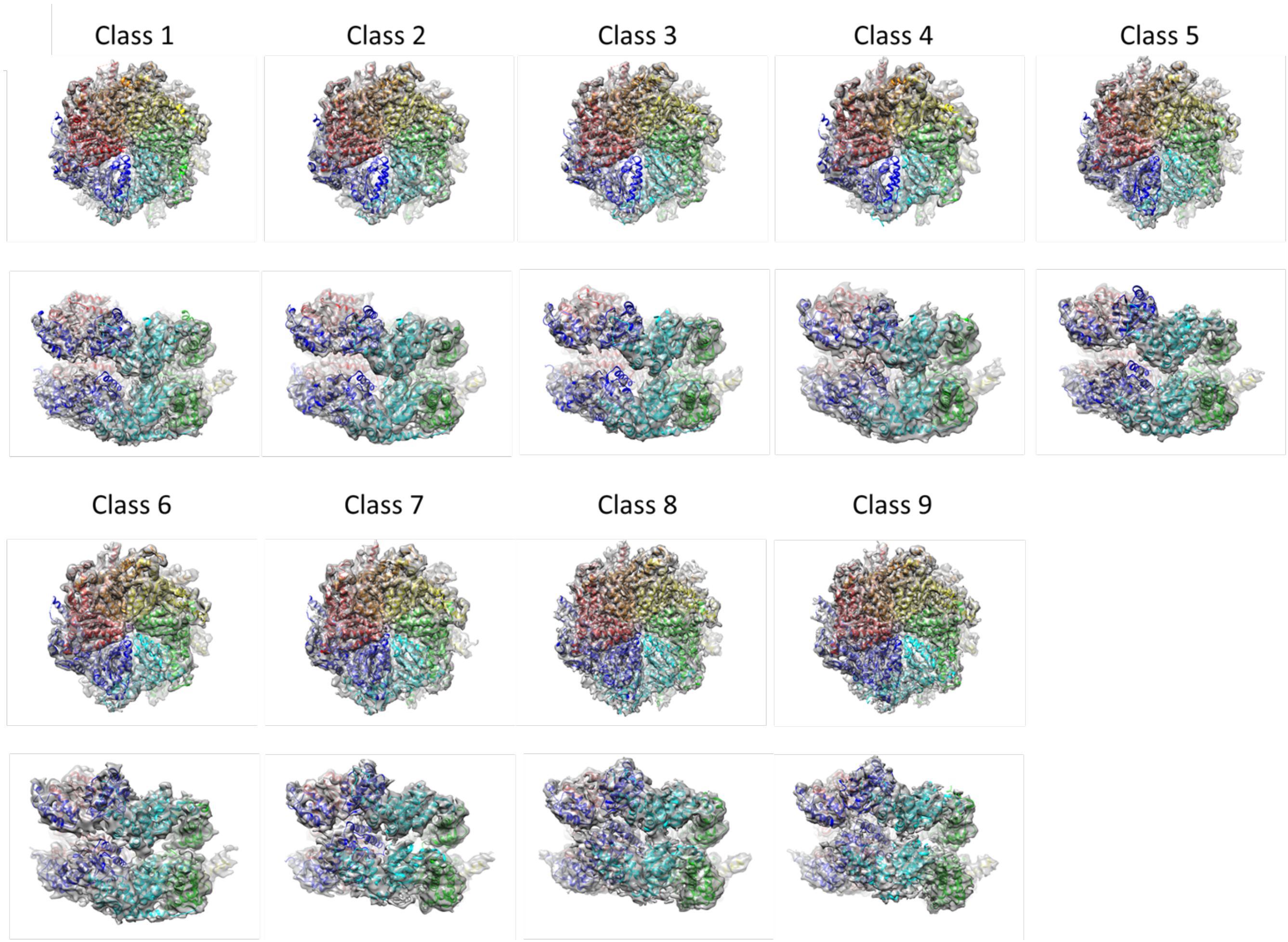


**Supplemental Figure 3. 3D Variability Analysis leads to 9 distinct classes with variable positioning of mobile subunits.** Fits of class 1-9 models into their respective maps at 4σ.

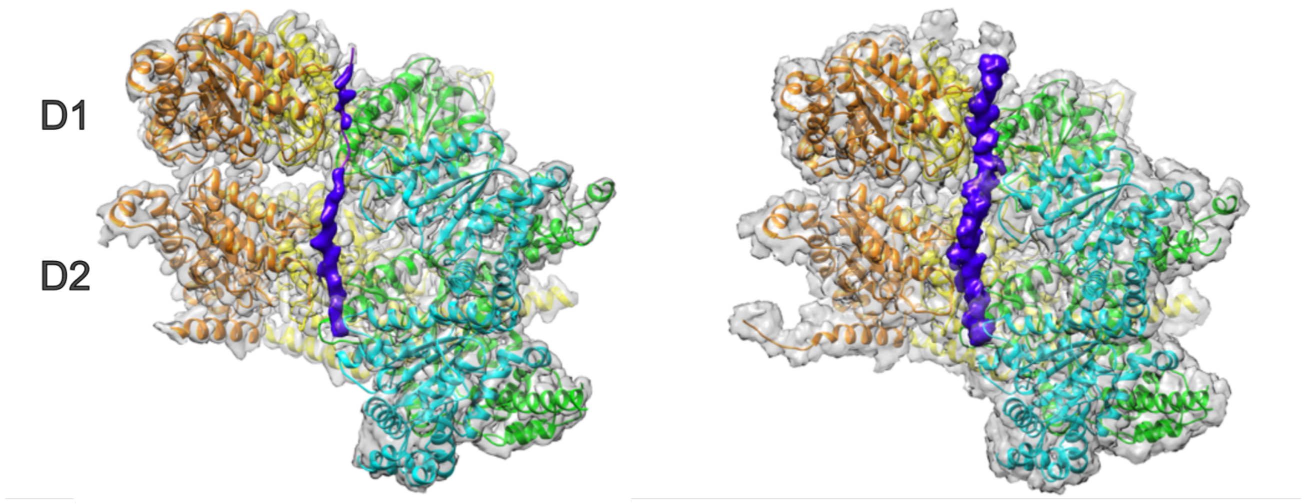


**Supplemental Figure 4. Substrate density at high and low thresholds.** Substrate density in purple with density displayed at 6σ (left) and at 2.5σ (right).

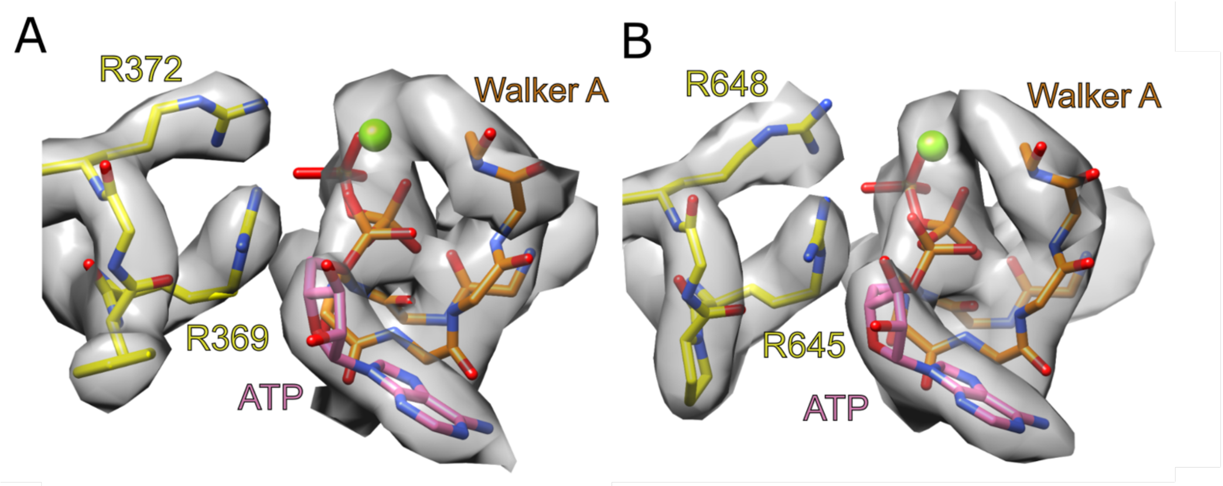


**Supplemental Figure 5. Coordination between arginine fingers and ATP.** (A) D1 nucleotide binding pocket at the BC interface with arginine fingers (R369, R372) from subunit C. Threshold at 7.5σ. (B) D2 nucleotide binding pocket at the BC interface with arginine fingers (R645, R648). Threshold at 8.75σ.

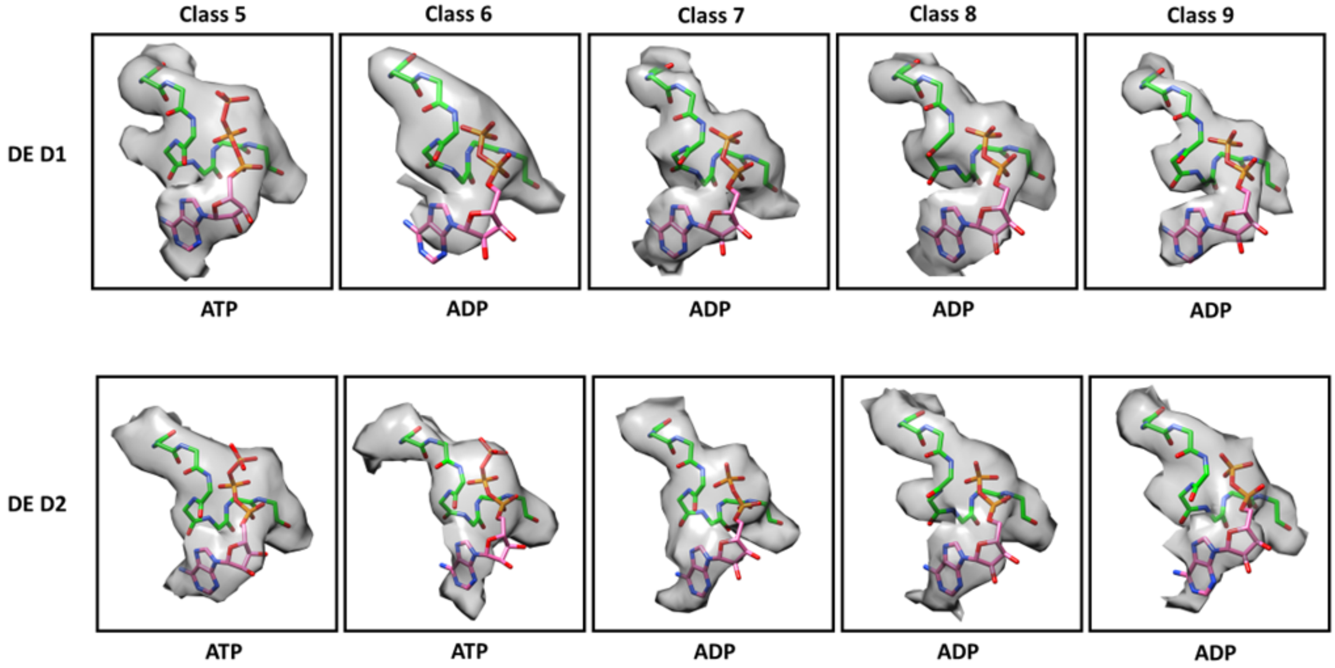


**Supplemental Figure 6. Model and density of nucleotides and the Walker A motif at the DE interface.** D1 (top) and D2 (bottom). Density displayed at 7$\sigma$. Walker A in green, nucleotide in pink.

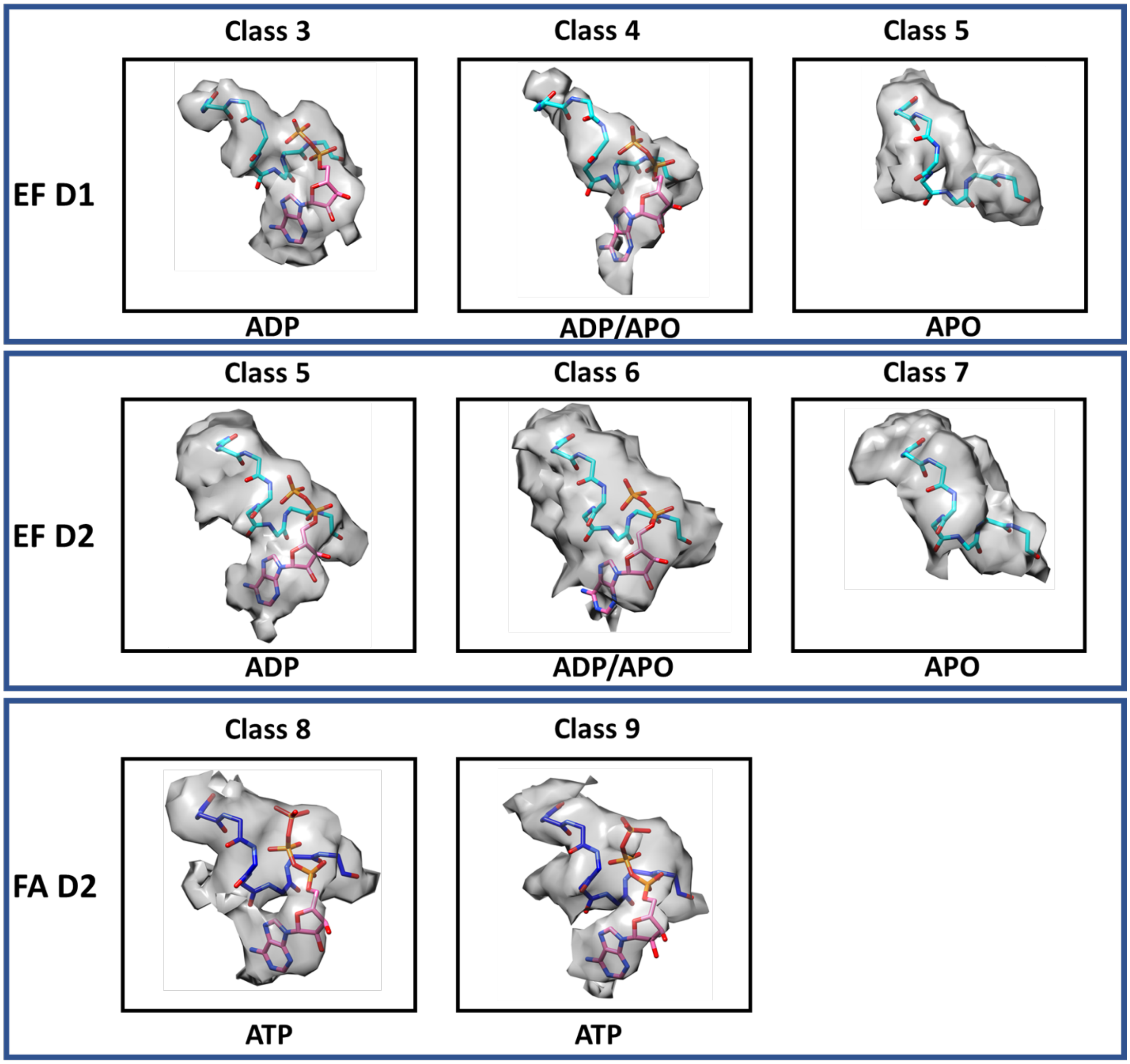


**Supplemental Figure 7. Model and density of nucleotides and the Walker A motif at EF and FA interfaces**. Density displayed at 5$\sigma$.

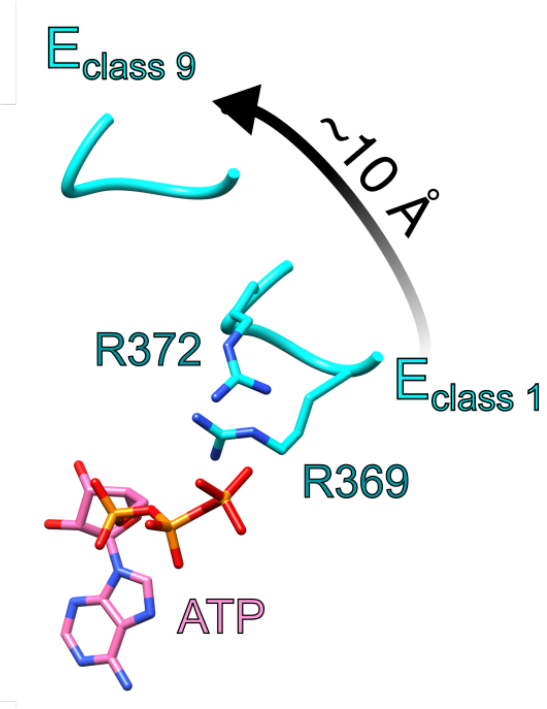


**Supplemental Figure 8. Displacement of arginine fingers during subunit movement.** Trajectory of arginine finger movement as subunit E of D1 moves away from the nucleotide binding pocket at the DE interface between classes 1-9. A similar range of motion is observed in subunit E of D2.

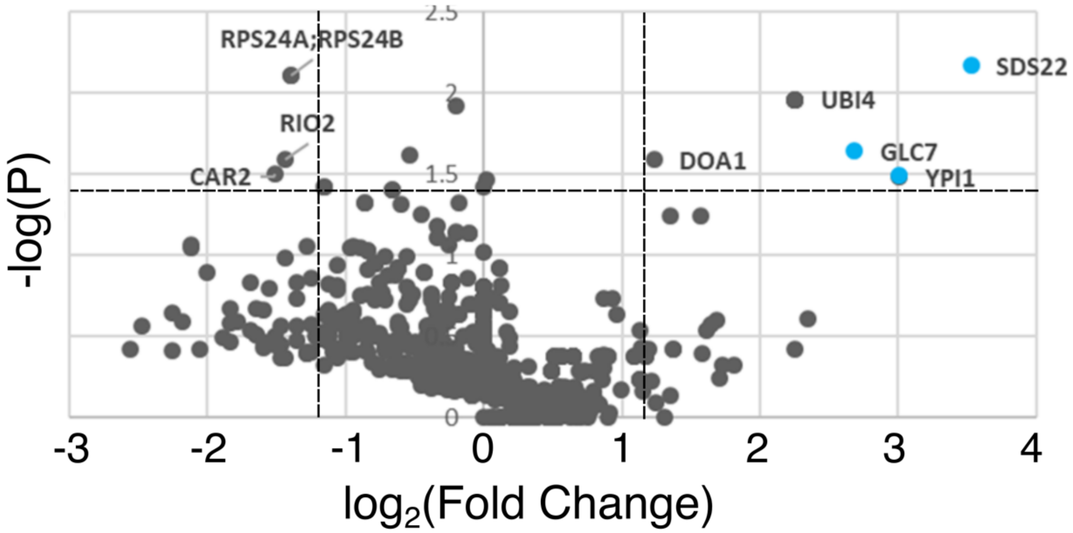


| Gene | FoldChange (Log 2) | P value (-Log) | Peptide sequences | Coverage |
|---|---|---|---|---|
| SDS22 | 3.5 | 2.2 | 28 | 82% |
| YPI1 | 3.0 | 1.5 | 12 | 52% |
| GLC7 | 2.7 | 1.6 | 17 | 52% |
| UBI4 | 2.3 | 2.0 | 15 | 90% |
| DOA1 | 1.2 | 1.6 | 57 | 77% |
| DBP2 | -1.2 | 1.4 | 22 | 50% |
| RPS24A | -1.4 | 2.1 | 4 | 29% |
| RIO2 | -1.4 | 1.6 | 6 | 19% |
| CAR2 | -1.5 | 1.5 | 73 | 80% |

**Supplemental Figure 9. Mass spectrometry proteomics of ATP elutions from Shp1 co-IPs**. Top, Volcano plot of detected proteins from Shp1-FLAG co-IPs eluted with ATP over ADP•BeF$_x$ controls. Bottom, list of proteins from A with –log(P) ≥ 1.4 and log$_2$FC ≥|1.2|.

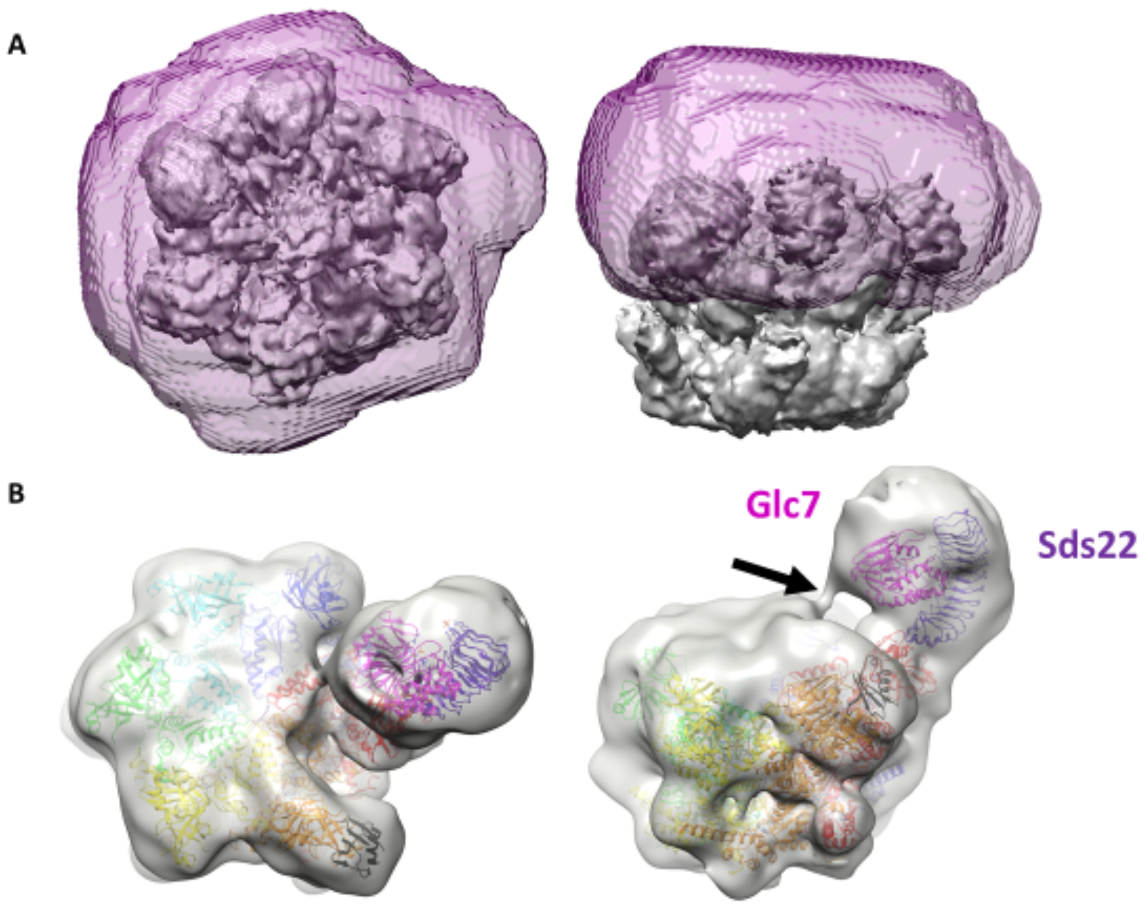


**Supplemental Figure 10. Image processing of PP1-containing class (class P)**. (A) Mask used for 3DVA of Cdc48 N-terminal domains. (B) Map at 1σ reveals connecting density between Glc7-Sds22 to the Cdc48 central pore.

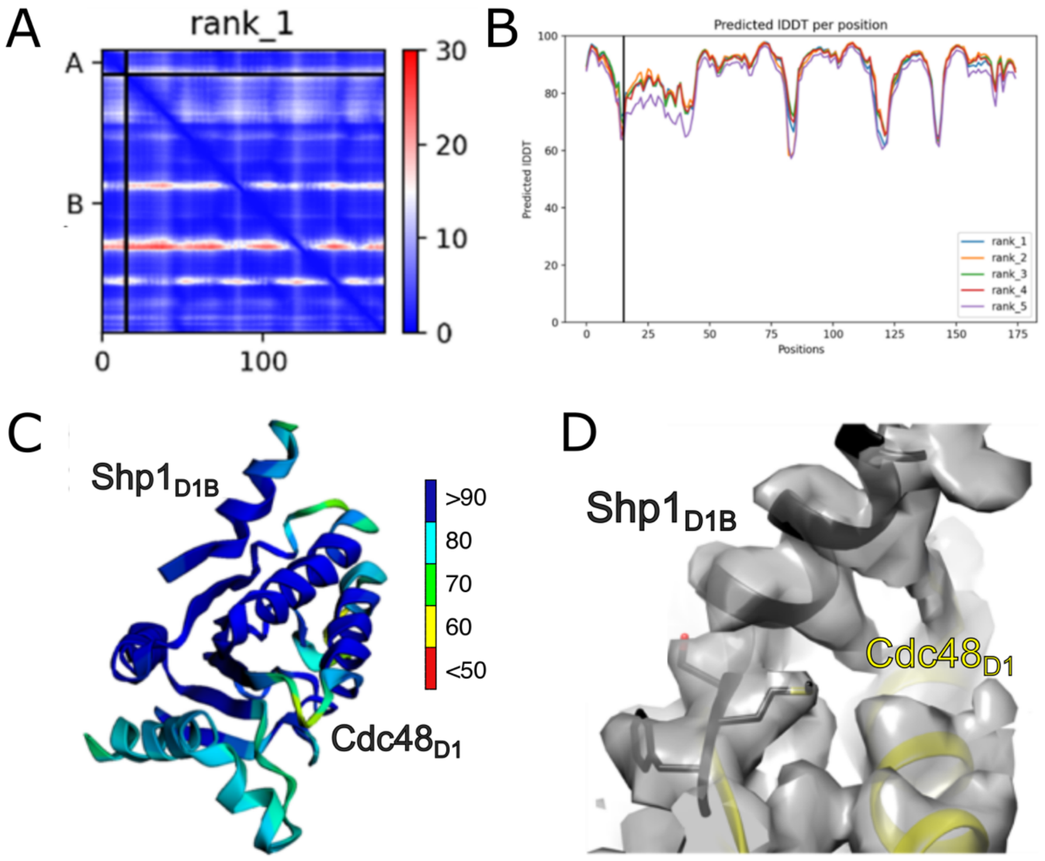


**Supplemental Figure 11. ColabFold prediction of Shp1-Cdc48 interaction.** (A) Predicted aligned error map for ColabFold prediction. Chain A, Shp1 D1B motif (residues 125-139); Chain B, Cdc48 D1 domain (residues 219-379). (B) Predicted local distance difference test (lDDT) scores for Shp1 D1B and Cdc48 D1. (C) Confidence scores for positions of predicted Shp1 D1B interactions with Cdc48 D1. (D) ColabFold predicted model of Shp1 D1B fitted as a rigid body into Cdc48 D1 of subunit C within the consensus reconstruction.

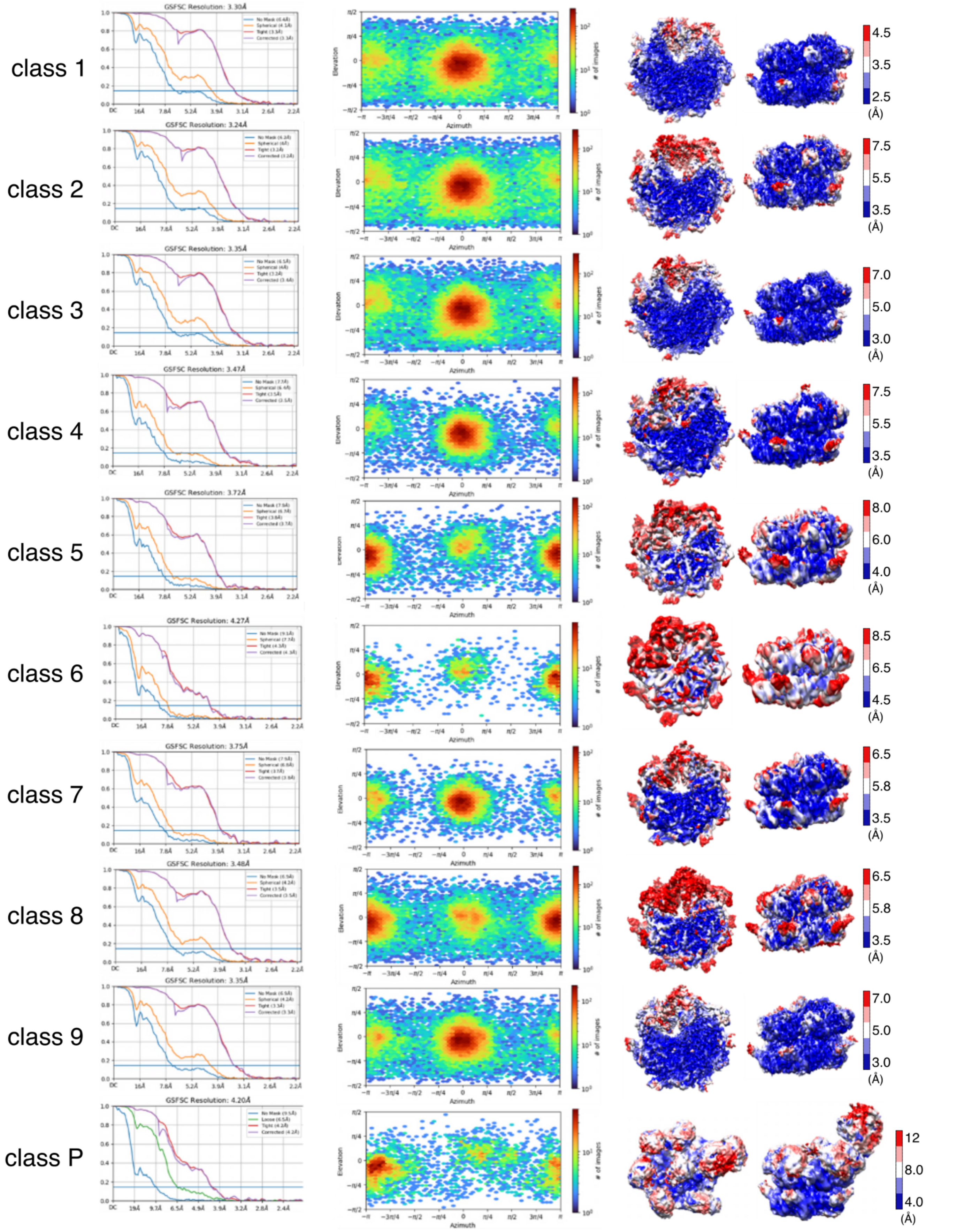


**Supplemental Figure 12. Cryo-EM reconstruction validation.** Gold standard FSC (left), orientation distribution plot (middle), and local resolution heat map (right).

**Supplemental Table 1.** Assigned nucleotide states for each subunit in classes 1-9 in D1 (left) and D2 (right).

## D1

| Class number | AB | BC | CD | DE | EF | FA |
|---|---|---|---|---|---|---|
| 9 | ATP | ATP | ATP | ADP | APO | APO |
| 8 | ATP | ATP | ATP | ADP | APO | APO |
| 7 | ATP | ATP | ATP | ADP | APO | APO |
| 6 | ATP | ATP | ATP | ADP/ATP | APO | APO |
| 5 | ATP | ATP | ATP | ATP | APO | APO |
| 4 | ATP | ATP | ATP | ATP | ADP/APO | APO |
| 3 | ATP | ATP | ATP | ATP | ADP | APO |
| 2 | ATP | ATP | ATP | ATP | ADP | APO |
| 1 | ATP | ATP | ATP | ATP | ADP | APO |

## D2

| Class number | AB | BC | CD | DE | EF | FA |
|---|---|---|---|---|---|---|
| 9 | ATP | ATP | ATP | ADP | APO | ATP |
| 8 | ATP | ATP | ATP | ADP | APO | ATP |
| 7 | ATP | ATP | ATP | ADP/ATP | APO | APO |
| 6 | ATP | ATP | ATP | ATP | ADP/APO | APO |
| 5 | ATP | ATP | ATP | ATP | ADP | APO |
| 4 | ATP | ATP | ATP | ATP | ADP | APO |
| 3 | ATP | ATP | ATP | ATP | ADP | APO |
| 2 | ATP | ATP | ATP | ATP | ADP | APO |
| 1 | ATP | ATP | ATP | ATP | ADP | APO |

### Supplemental Table 2. Validation Statistics (1/2)

| | Consensus | Class 1 | Class 2 | Class 3 | Class 4 |
|---|---|---|---|---|---|
| **Collection and Data Processing** | | | | | |
| Magnification | 81000x | 81000x | 81000x | 81000x | 81000x |
| Defocus Range (μm) | .5-5 | .5-5 | .5-5 | .5-5 | .5-5 |
| Number of micrographs | 5847 | 5847 | 5847 | 5847 | 5847 |
| Initial particle images | 4,690,491 | 4,690,491 | 4,690,491 | 4,690,491 | 4,690,491 |
| Symmetry imposed | C1 | C1 | C1 | C1 | C1 |
| Final Particle images | 325,377 | 41,227 | 45,203 | 42,875 | 22,656 |
| Map Resolution FSC 0.143 (Å) | | | | | |
| Masked | 2.9 | 3.3 | 3.2 | 3.4 | 3.5 |
| Unmasked | 3.6 | 6.4 | 6.2 | 6.5 | 7.7 |
| **Validation** | | | | | |
| Starting model used (PDB code) | 6OPC | 6OPC | 6OPC | 6OPC | 6OPC |
| Map Correlation coefficient | 0.75 | 0.74 | 0.73 | 0.74 | 0.71 |
| Atoms | 43698 | 43396 | 44137 | 45305 | 39205 |
| Protein residues | 3246 | 3174 | 3187 | 3215 | 3158 |
| Bonds (RMSD) | | | | | |
| Length (Å) | 0.003 (0) | 0.003 (0) | 0.003 (0) | 0.004 (0) | 0.004 (0) |
| Angles (°) | 0.617 (0) | 0.673 (2) | 0.696 (0) | 0.682 (2) | 0.671 (0) |
| MolProbity score | 1.37 | 1.55 | 1.65 | 1.63 | 1.53 |
| Clash score | 2.41 | 4.3 | 5.54 | 5.51 | 3.63 |
| Ramachandran plot (%) | | | | | |
| Outliers | 0 | 0 | 0 | 0.03 | 0 |
| Allowed | 4.96 | 4.89 | 4.99 | 4.72 | 5.47 |
| Favored | 95.04 | 95.11 | 95.01 | 95.25 | 94.53 |
| Rotamer outliers (%) | 0 | 0 | 0 | 0 | 0 |
| Cß outliers (%) | 0 | 0 | 0 | 0 | 0 |
| CaBLAM outliers (%) | 3.84 | 3.65 | 3.43 | 3.55 | 3.64 |

**Supplemental Table 2. Validation Statistics, continued (2/2)**

| | Class 5 | Class 6 | Class 7 | Class 8 | Class 9 |
|---|---|---|---|---|---|
| **Collection and Data Processing** | | | | | |
| Magnification | 81000x | 81000x | 81000x | 81000x | 81000x |
| Defocus Range (µm) | .5-5 | .5-5 | .5-5 | .5-5 | .5-5 |
| Number of micrographs | 5847 | 5847 | 5847 | 5847 | 5847 |
| Initial particle images | 4,690,491 | 4,690,491 | 4,690,491 | 4,690,491 | 4,690,491 |
| Symmetry imposed | C1 | C1 | C1 | C1 | C1 |
| Final Particle images | 16,846 | 6,164 | 14,639 | 35,398 | 47,580 |
| Map Resolution FSC 0.143 (Å) | | | | | |
| Masked | 3.7 | 4.3 | 3.8 | 3.5 | 3.4 |
| Unmasked | 7.9 | 9.1 | 7.9 | 6.9 | 6.9 |
| **Validation** | | | | | |
| Starting model used (PDB code) | 6OPC | 6OPC | 6OPC | 6OPC | 6OPC |
| Map Correlation coefficient | 0.73 | 0.73 | 0.71 | 0.72 | 0.75 |
| Atoms | 44496 | 43067 | 43622 | 43497 | 42357 |
| Protein residues | 3229 | 3244 | 3204 | 3194 | 3166 |
| Bonds (RMSD) | | | | | |
| Length (Å) | 0.003 (0) | 0.003 (0) | 0.003 (0) | 0.006 (0) | 0.003 (0) |
| Angles (°) | 0.670 (0) | 0.710 (3) | 0.731 (4) | 0.773 (2) | 0.638 (0) |
| MolProbity score | 1.55 | 1.62 | 1.7 | 1.62 | 1.49 |
| Clash score | 4.62 | 5.58 | 6.76 | 4.25 | 4.43 |
| Ramachandran plot (%) | | | | | |
| Outliers | 0 | 0 | 0.03 | 0 | 0 |
| Allowed | 4.51 | 4.56 | 4.68 | 6.23 | 3.98 |
| Favored | 95.49 | 95.44 | 95.29 | 93.77 | 96.02 |
| Rotamer outliers (%) | 0 | 0 | 0 | 0 | 0 |
| Cß outliers (%) | 0 | 0 | 0 | 0 | 0 |
| CaBLAM outliers (%) | 3.53 | 3.82 | 3.31 | 3.78 | 2.75 |
